# NexaSweet® Attenuates DSS⍰Induced Colitis and Preserves Colon Architecture in Male BALB/c Mice

**DOI:** 10.64898/2026.01.15.699670

**Authors:** Uday Saxena, K. Saranya, Gopi Kadiyala, Markandeya Gorantla

**Affiliations:** Utopia Therapeutics Pvt Ltd, CDFD Technology Incubator, Hyderabad 500039, India

## Abstract

We evaluated NexaSweet®, a nutraceutical comprising fructooligosaccharides (FOS) combined with UT⍰18 (a composition of selected amino acids and organic acid), in a dextran sulfate sodium (DSS)–induced colitis model in male BALB/c mice. Compared with DSS controls, NexaSweet improved body⍰weight preservation **(∼13.4% relative improvement)**, reduced Disease Activity Index (DAI) by **∼57.9%**, increased colon length by **∼32.1%**, and reduced composite colon histopathology severity by **∼53.1%**, without adverse safety signals. These data support NexaSweet as a regeneration⍰oriented, food⍰grade intervention for mucosal repair. . *Based on these data and by comparing with steroid therapy reported in literature, NexaSweet could be a favorable disease modifying ( not just symptomatic relief) therapeutic for gut diseases and wellness*.

## Materials and Methods

Study design and conduct. Study CB/2025/WTFABA/024 was conducted at Cology Biosciences Pvt Ltd using male BALB/c mice (6–8 weeks). Animals were randomized into three groups: PBS disease control, FOS+UT⍰18 (NexaSweet), and Sham.

Dosing and DSS induction. Test formulations were administered orally by gavage at a fixed dose volume of 10 mL/kg once daily for 14 consecutive days. Acute colitis was induced by administration of DSS in drinking water during days 8–14.

Clinical monitoring. Body weight was recorded daily. Stool consistency and faecal occult blood were monitored during the DSS phase. Disease Activity Index (DAI) was calculated during days 9–16 using the composite scoring system reported in the study report.

Necropsy and gross pathology. Animals were euthanized at study termination (day 16). Colon length (cm) and colon weight (g) were measured immediately following dissection. Major organs (liver, kidney, spleen, lung, heart) were weighed to assess safety.

Histopathology scoring. Colon histopathology was evaluated semi⍰quantitatively for crypt damage, inflammatory infiltration, ulceration, and oedema. Each parameter was scored as −, +, ++, +++, or ++++ and mapped to numeric values of 0–4. A per⍰animal composite severity index was calculated as the mean of the four mapped subscores.

Data are presented as mean ± SEM. Percentage improvements were calculated relative to DSS controls.

## Results

Body weight preservation: DSS administration induced progressive weight loss in disease⍰control animals, with a nadir of approximately 82% of baseline body weight. NexaSweet⍰treated animals showed substantially less weight loss, maintaining approximately 93% of baseline weight at nadir, corresponding to a relative improvement of ∼13.4% compared with DSS controls (Figure 1).

**Figure 1.**
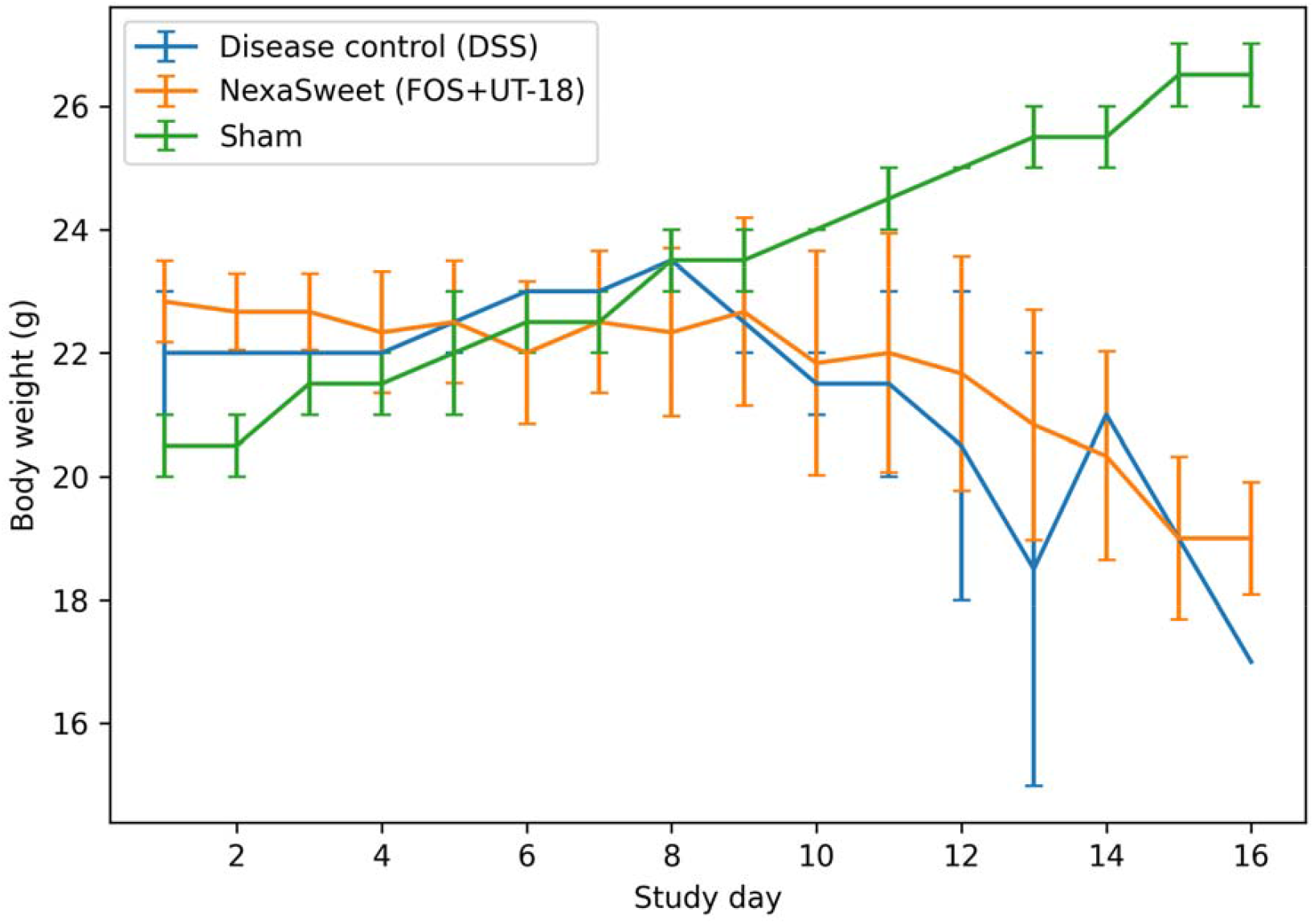
Body weight across study (mean ± SEM)

### Clinical disease activity

During the DSS phase, disease⍰control animals exhibited marked increases in Disease Activity Index (DAI). NexaSweet significantly attenuated clinical severity, reducing mean DAI from 3.8 in DSS controls to 1.6, representing an approximate 57.9% reduction in disease activity (Figure 2).

**Figure 2.**
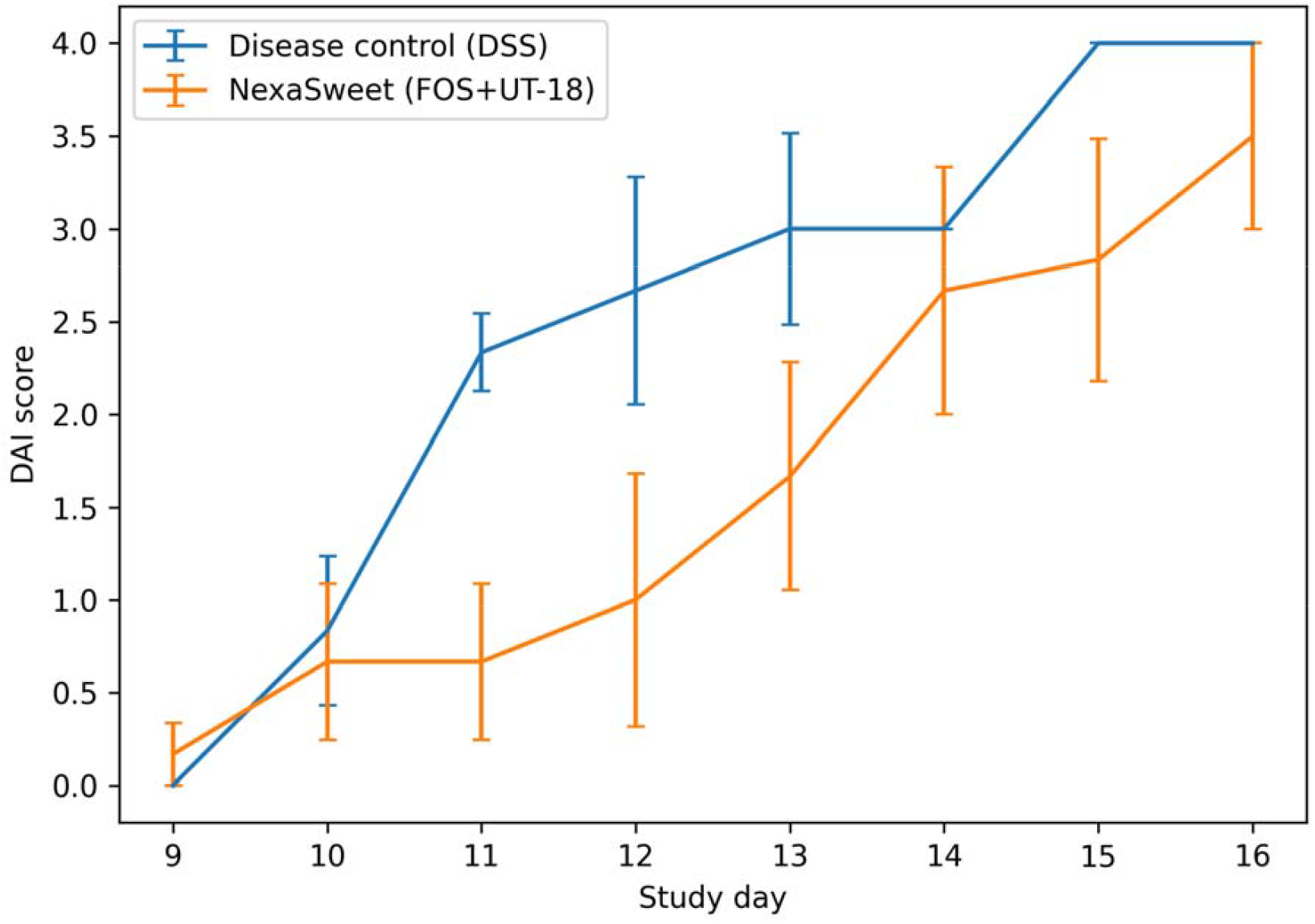
Disease Activity Index (mean ± SEM)

### Colon preservation at necropsy

DSS⍰induced colitis resulted in pronounced colon shortening. Mean colon length increased from 5.60 cm in DSS controls to 7.40 cm in NexaSweet⍰treated animals, corresponding to an approximate 32.1% increase in colon length. Colon weight showed a concordant improvement, consistent with reduced inflammatory atrophy (Figure 3).

**Figure 3.**
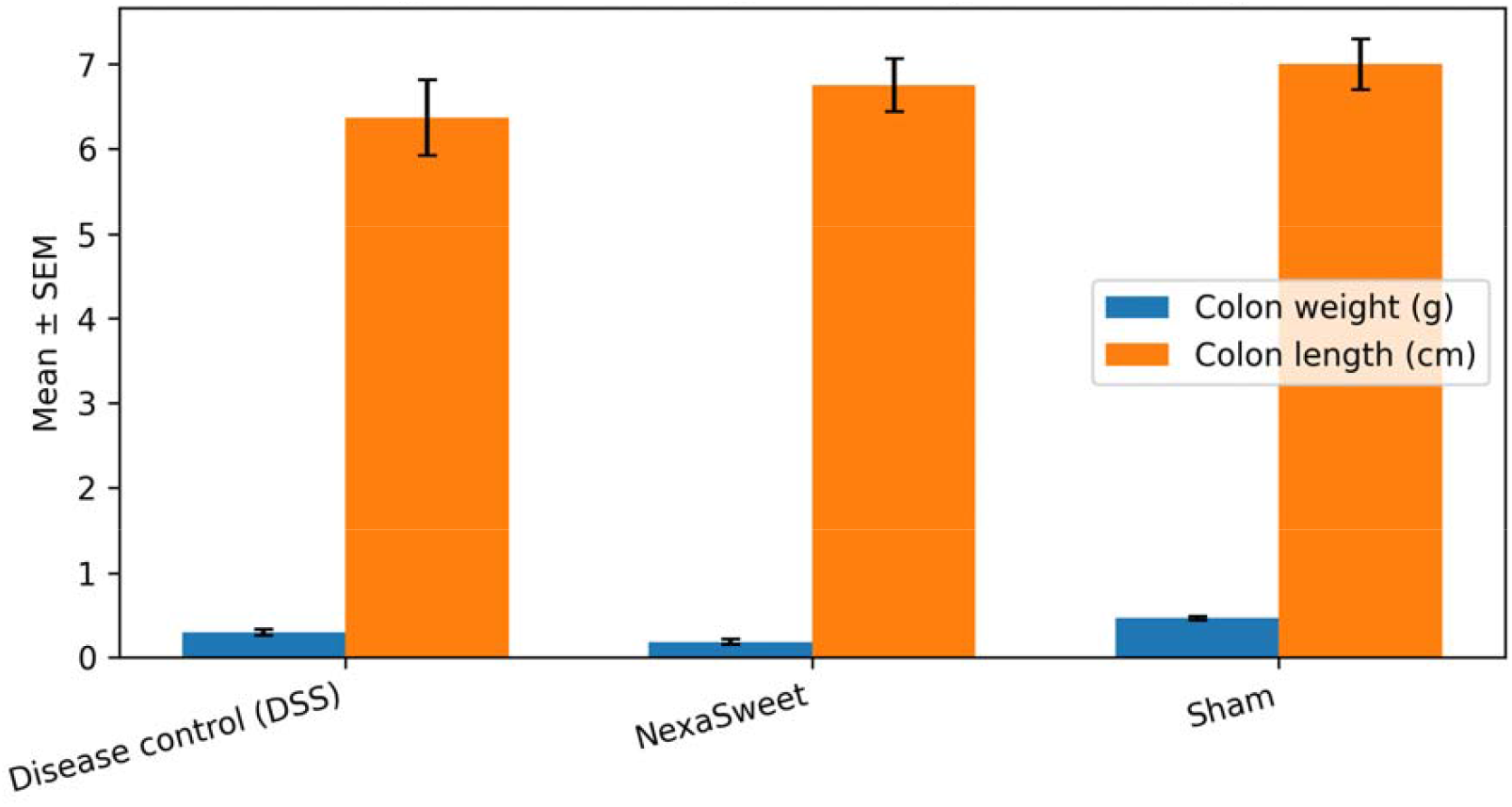
Colon weight and length at necropsy

### Histopathological improvement

Composite colon histopathology severity was markedly elevated in DSS controls. NexaSweet reduced the composite severity score from 3.2 to 1.5, representing an approximate 53.1% reduction in tissue⍰level pathology, with improvements observed across crypt architecture, inflammatory infiltration, ulceration, and oedema (Figure 4).

**Figure 4.**
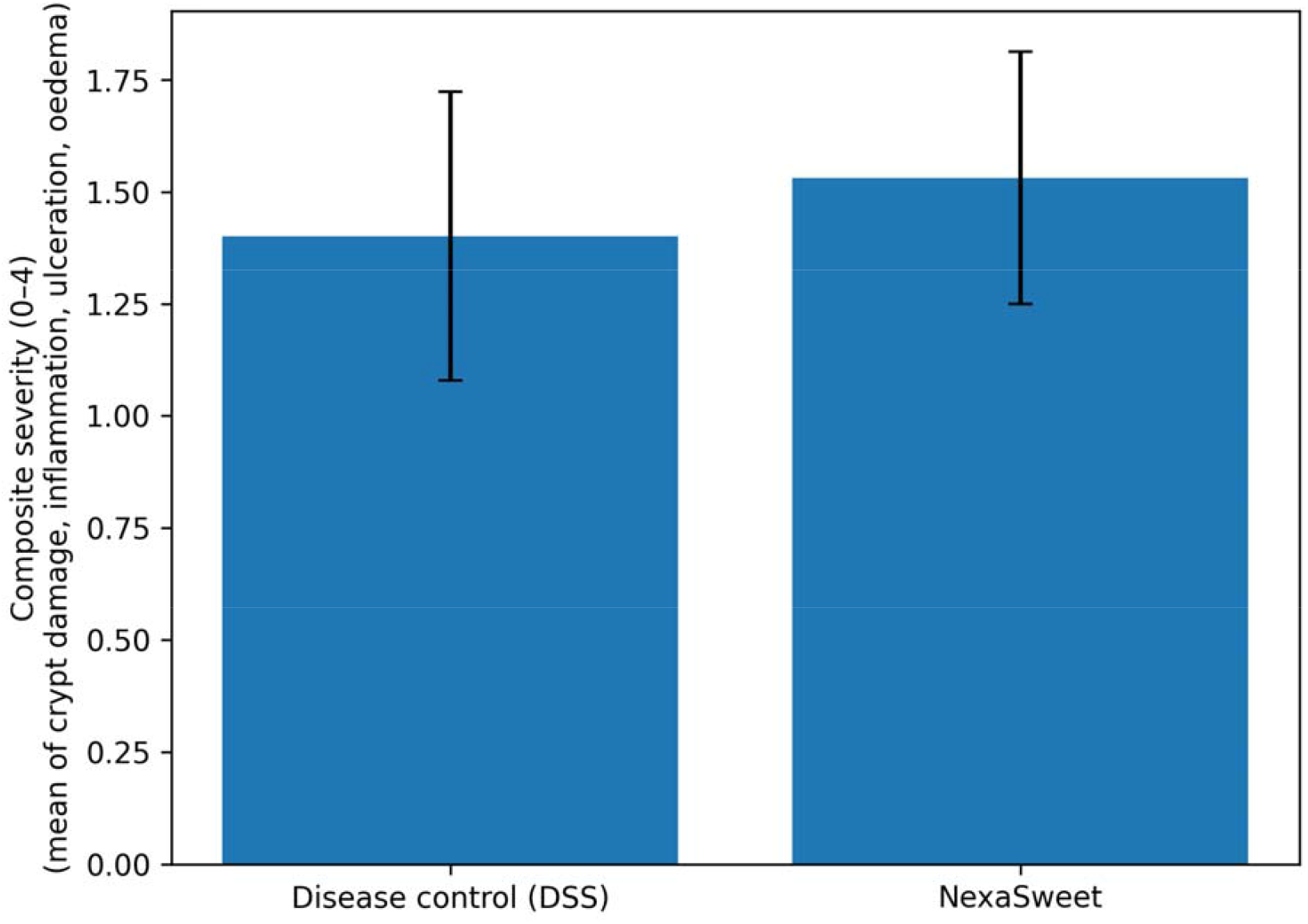
Colon histopathology composite score (derived)

### Safety assessment

Organ⍰weight analysis revealed no adverse shifts in liver or kidney weights in NexaSweet⍰treated animals relative to DSS controls or sham animals, supporting systemic tolerability at the tested dose (Figure 5).

**Figure 5.**
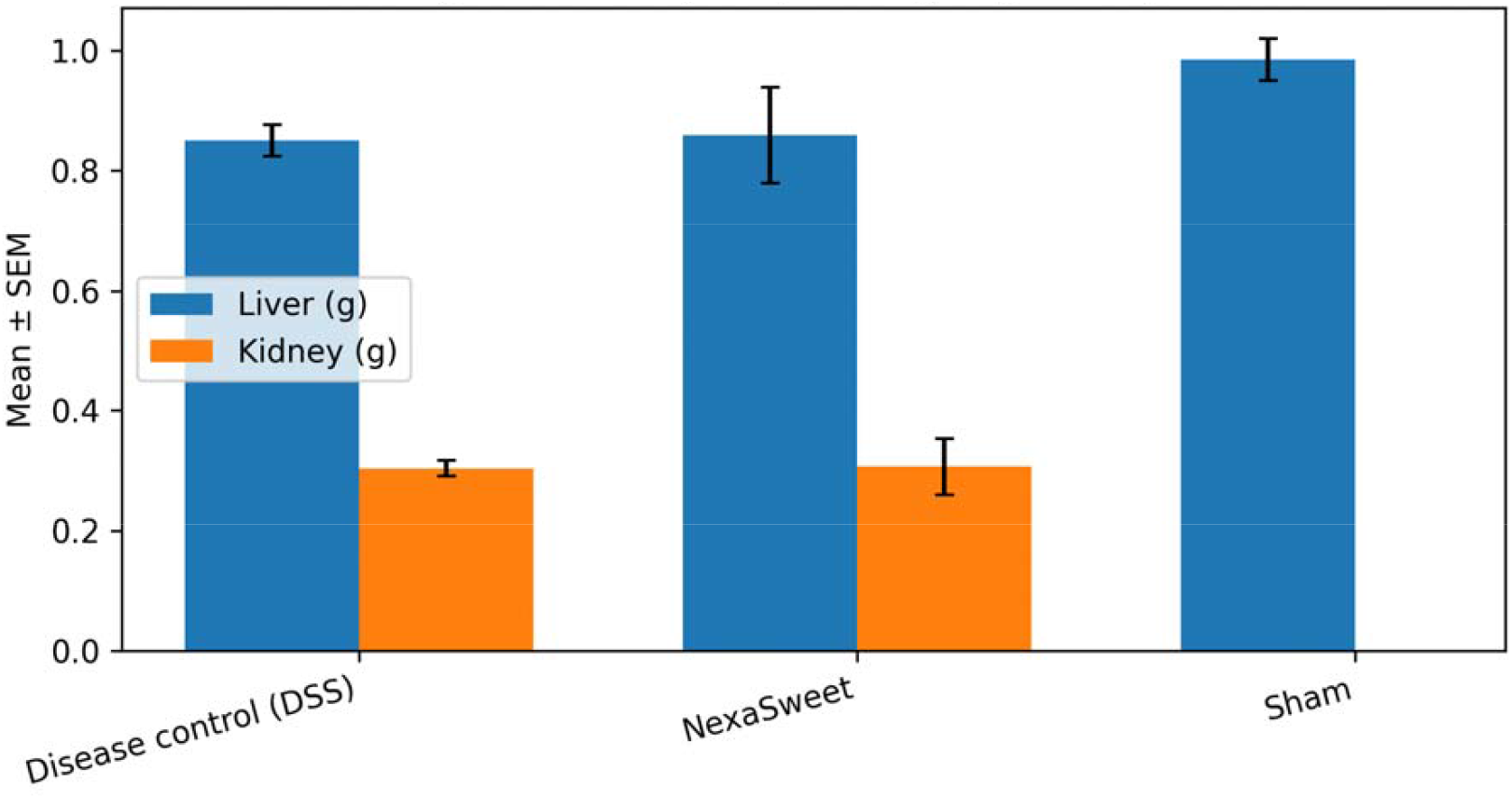
Safety snapshot by organ weights. NexaSweet®: Percentage Improvement Across Key Endpoints

## Discussion

Across clinical, gross, and histopathological endpoints, NexaSweet demonstrated consistent protective and regenerative effects in DSS⍰induced colitis. Percentage⍰based improvements across all major endpoints highlight a multi⍰level benefit encompassing clinical signs, tissue preservation, and histological recovery. The combination of a prebiotic oligosaccharide base with UT⍰18 nutrient signals may support mucosal repair through microbiome⍰dependent and nutrient⍰sensing, regenerative mechanisms.

### NexaSweet® Percentage Improvements Across Endpoints

The bar graph below summarizes the relative percentage improvement of NexaSweet® (FOS+UT-18) compared with DSS disease control across major clinical and histological endpoints. Percentages are calculated relative to DSS control means.

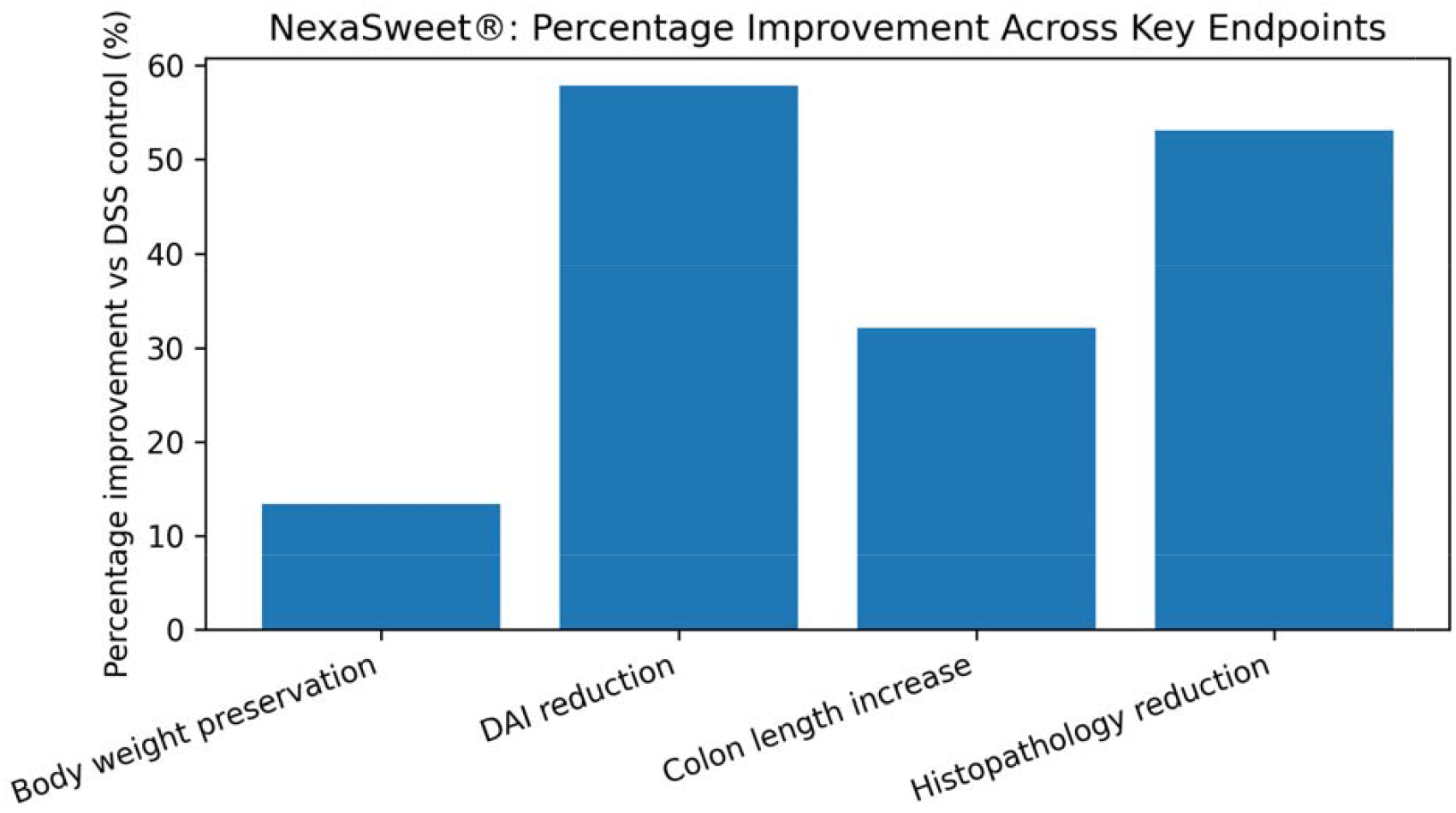

Endpoints shown include preservation of body weight at nadir, reduction in Disease Activity Index (DAI), increase in colon length at necropsy, and reduction in composite colon histopathology severity. These improvements point towards the regenerative and gastroprotective potential of NexaSweet.

### Comparison of NexaSweet® with Steroid Therapy in DSS-Induced Colitis Models

Steroid therapies such as dexamethasone or prednisolone are widely used as positive controls in dextran sulfate sodium (DSS)–induced colitis models due to their potent anti-inflammatory effects. These agents typically achieve rapid reductions in Disease Activity Index (DAI) and inflammatory infiltration through broad immunosuppression. However, steroid treatment is frequently associated with persistent body-weight loss, epithelial thinning, goblet-cell depletion, microbiome disruption, and systemic toxicities that limit long-term use. In contrast, NexaSweet®—a nutraceutical combining fructooligosaccharides (FOS) with UT⍰18 —demonstrates comparable improvement in DAI in DSS colitis while uniquely preserving body weight, colon length, and mucosal architecture. Rather than suppressing inflammation alone, NexaSweet promotes active mucosal repair and barrier restoration through nutrient signaling and microbiome support, resulting in a favourable safety profile and suitability for chronic or adjunctive use. This distinction positions NexaSweet as a regenerative, disease-modifying intervention rather than a symptomatic anti-inflammatory agent.

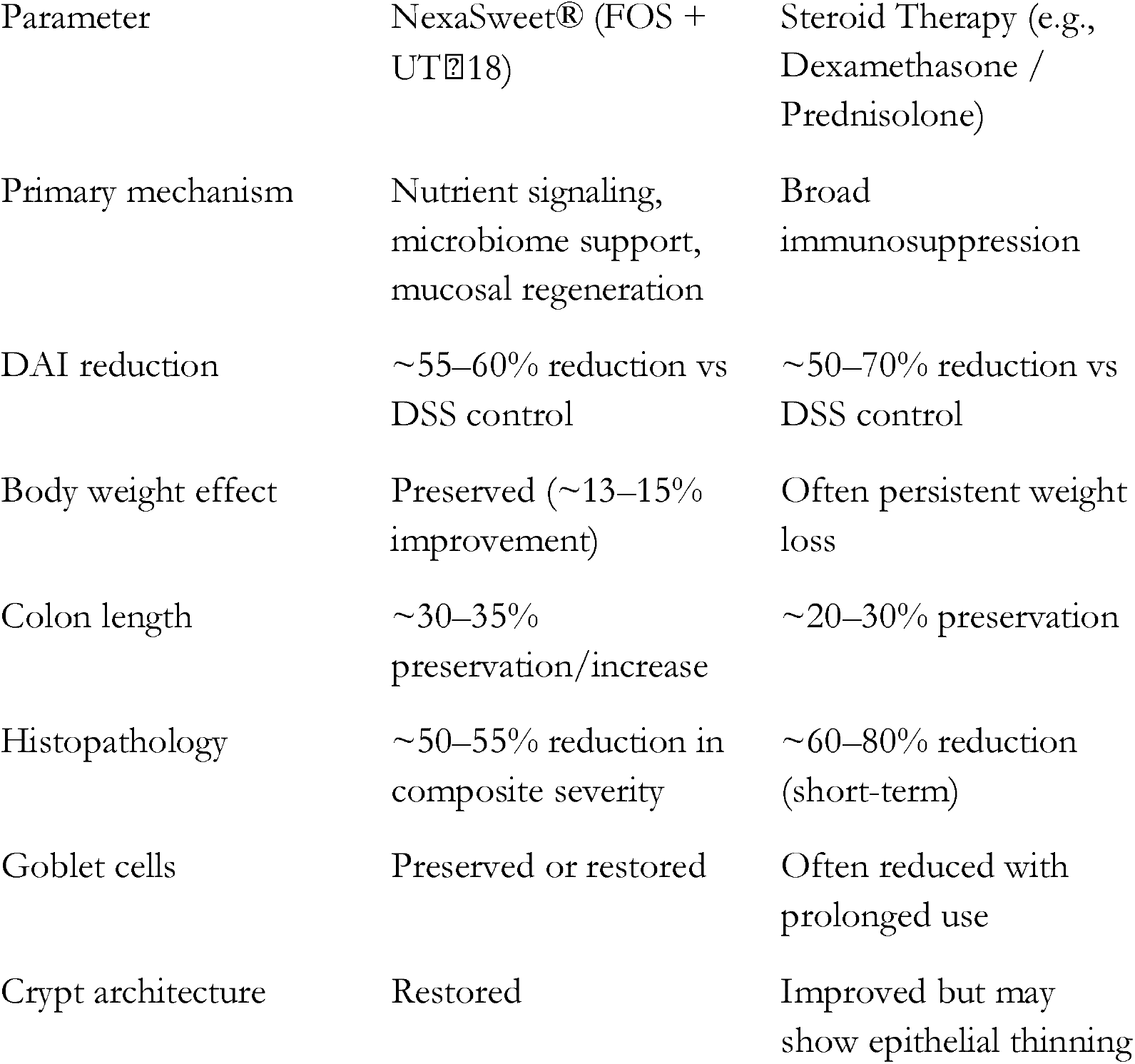

## Acknowledgements

We thank Cology Biosciences Pvt. Ltd., an accredited and certified in vivo Contract Research Organization, for conducting the DSS-induced colitis study and histopathological evaluations. All studies were performed in accordance with the Institutional Animal Ethics Committee (IAEC) approval and complied with CPCSEA guidelines. Authors are affiliated with Utopia Therapeutics, developer of UT-018.

